# Calcium and strontium contractile activation properties of single skinned skeletal muscle fibres from elderly women 66-90 years of age

**DOI:** 10.1101/2021.11.08.466855

**Authors:** Susan M. Ronaldson, D. George Stephenson, Stewart I. Head

**Affiliations:** Sydney Nursing School, The University of Sydney, Australia, 2050; Department of Zoology, La Trobe University, Melbourne, Australia, 3086; School of Medicine, Western Sydney University, Sydney, Australia, 2751

**Keywords:** human aging, calcium, skeletal muscle contraction, skeletal muscle fibre types

## Abstract

The single skinned muscle fibre technique was used to investigate Ca^2+^- and Sr^2+^- activation properties of skeletal muscle fibres from elderly women (66-90 years). Muscle biopsies were obtained from the vastus lateralis muscle. Three populations of muscle fibres were identified according to their specific Sr^2+^- activation properties: slow-twitch (type I), fast-twitch (type II) and hybrid (type I/II) fibres. All three fibre types were sampled from the biopsies of 66 to 72 years old women, but the muscle biopsies of women older than 80 years yielded only slow-twitch (type I) fibres. The proportion of hybrid fibres in the vastus lateralis muscle of women of circa 70 years of age (24%) was several-fold greater than in the same muscle of adults (<10%), suggesting that muscle remodelling occurs around this age. There were no differences between the Ca^2+^- and Sr^2+^- activation properties of slow-twitch fibres from the two groups of elderly women, but there were differences compared with muscle fibres from adults with respect to sensitivity to Ca^2+^, steepness of the activation curves, and characteristics of the fibre-type dependent phenomenon of spontaneous force oscillations (SOMO) occurring at submaximal levels of activation. The maximal Ca^2+^ activated specific force from all the fibres collected from the seven old women use in the present study was significantly lower by 20% than in the same muscle of adults. Taken together these results show there are qualitative and quantitative changes in the activation properties of the contractile apparatus of muscle fibres from the vastus lateralis muscle of women with advancing age, and that these changes need to be considered when explaining observed changes in women’s mobility with aging.

## INTRODUCTION

There have been a large number of reports of a loss of skeletal muscle mass and force generation capacity as a result of the aging process (Aniansson et al., 1980; Aniansson et al., 1983; Grimby et al., 1984; Brooks and Faulkner, 1988, 1991; Edstrom and Larsson, 1987; Jubrias et al., 1997; Lexell et al., 1983; Lexell et al., 1988; Sipilä and Suominen, 1991 Lamboley et al. 2015). However, it is not yet clear to what extent this reduction in force is due to an intrinsic loss of force generating capacity of the contractile proteins in a particular type of fibre, to the change in the fibre type composition, or to a reduction of the cross-sectional area of the muscle.

Several studies have shown a decrease or selective atrophy of specific muscle fibre types. In particular, it appears that fast-twitch muscle fibres are more prone to atrophy or damage with age (Aniansson et al., 1980; Larsson et al., 1978; Lexell et al., 1988; Lexell, 1995). Brocca et.al. (2017) found there was a switch from fast-twitch to slow-twitch fibres in the leg muscles from older men. While Meznaric et.al. (2018) found that there was a shift in the myosin heavy chain phenotype to the slower myosin heavy chain phenotype in the neck muscles of ageing males. They attributed this shift in the fibre type portion to a greater loss of fast-twitch motor neurons during ageing.

In an earlier study using intact skeletal muscles Brooks and Faulkner (1988) found that the maximum force generated per cross-sectional area was less in whole fast twitch muscle from old mice. However, in a later study using single skinned muscle fibres they found that there was no difference in the maximum force per cross-sectional area generated by muscle fibres from young and old mice (Brooks and Faulkner, 1994). They did, however, note that the Ca^2+^ sensitivity of the contractile proteins was less in muscle fibres from old mice compared with adult mouse muscle fibres (Brooks and Faulkner, 1994). In a study from our laboratory on the effect of ageing on whole fast-twitch EDL muscles we showed that old muscles were stiffer and produce less specific force with a shift towards slow-twitch contractile properties as evidenced by a slowing of relaxation and increased resistance to fatigue. Interestingly, from the point of view of the present study, the effect of these age-related changes was greater in female mice compared to the male mice (Chan & Head 2010). Straight et.al. 2018 using single skinned fibres from ageing men and women found that the type I, type IIA and hybrid I/IIA fibres were less sensitive to Ca^2+^ when compared with young controls. In a study using single chemically skinned fibres from young and old men Ochala et al. 2007 found a reduced specific force in fibers from old men.

In the present study we investigated the calcium (Ca^2+^) - and strontium (Sr^2+^) - activation properties of freshly dissected single skeletal muscle fibres from elderly women (60-90 yrs) using the skinned muscle fibre technique. Strontium (Sr^2+^) was also used as an activator because it allows unequivocal differentiation between type I, type II and hybrid (type I/type II) fibres (O’Connell et al., 2004; Lamboley et al., 2013). This is because the sensitivity to Sr^2+^ ions depends on the troponin C (TnC) isoform (slow, fast or both slow and fast) expressed in the respective fibre, which in turn, is tightly correlated with the myosin heavy chain (MHC) expressed (MHCI or MHCII) in the fibre (Bortolotto et al., 2000, O’Connell et al., 2004; Lamboley et al., 2013). Moreover, the Sr^+^-activation curves of hybrid fibres permit direct estimation of the fraction of TnC isoforms present in the fibre (O’Connell et al., 2004).

## MATERIALS AND METHODS

### Preparation

Single muscle fibres were dissected from fresh muscle biopsies obtained from elderly women undergoing orthopaedic surgery for total hip replacement or repair of a fractured neck of femur. The women’s ages ranged from 66 to 90 years (n=7). Each woman’s activity profile was obtained from interview and medical records data prior to the scheduled surgery. Care was taken to select only women with an active lifestyle prior to surgery. Informed written consent was obtained from each woman and human ethics approval was obtained from both LaTrobe University and Austin Hospital, Melbourne, Australia, Human Research Ethics Committees.

The muscle sample was dissected by an orthopaedic surgeon from the vastus lateralis muscle in elderly women undergoing orthopaedic surgery. The muscle biopsy was obtained within 5-15 mins of the commencement of the surgery. The muscle biopsy was blotted thoroughly on Whatman’s filter paper (No 1) immediately upon its dissection from the vastus lateralis muscle to remove any excess interstitial fluid, then placed in a jar of cold paraffin oil at 2°C. The muscle biopsy was then placed in a thermos flask containing ice and transferred immediately to the laboratory.

The dissection of the muscle fibres from the muscle biopsy was generally commenced within 60 mins of its removal from the vastus lateralis muscle. The muscle biopsy was transferred from the cold paraffin into a Petri dish containing cold paraffin. Lynch et al. (1993) showed that this biopsy procedure had no effect on the activation characteristics of human skeletal muscle biopsy samples. Dissection of muscle fibres took place under oil. The muscle fibre was mechanically skinned (Stephenson and Williams, 1981) and tied at one end with silk thread and the other end was mounted between a pair of fixed forceps and attached to a force transducer (801 SensoNor Horten, Norway). Once a single muscle fibre had been dissected the remainder of the biopsy sample was stored under cold paraffin at 2°C for up to 6 hours during which further fibres were dissected out.

The length of the fibre was adjusted such that the preparation was just taut, and the diameter of the skinned muscle fibres was measured under oil. The sarcomere length (SL) of the skinned muscle fibres was then measured by laser diffraction (mean SL 2.71+/-0.04 μm; Stephenson and Williams, 1981).

### Solutions

Solutions were prepared according to standard procedures described by Stephenson and Williams (1981). The composition of the solutions is given in Table 1. Following mounting of the single skinned muscle fibre it was lowered into a relaxing solution (solution A Table 1) containing EGTA (50 mM) and allowed to equilibrate for 5 minutes. Before activation the fibre was immersed in a preactivating solution containing HDTA (solution D Table 1) to facilitate a rapid [Ca^2+^] rise (Moisescu & Thieleczek, 1978) when the fibre was placed in the Ca^2+^ activating solutions (solution B Table 1) after which it was returned to the relaxing solution (solution A Table 1). This procedure was repeated for activation of muscle fibres in Sr^2+^ solutions (solution C Table 1; Figure 1). All experiments were performed at room temperature (22 +/-1°C).

**Table 1.**
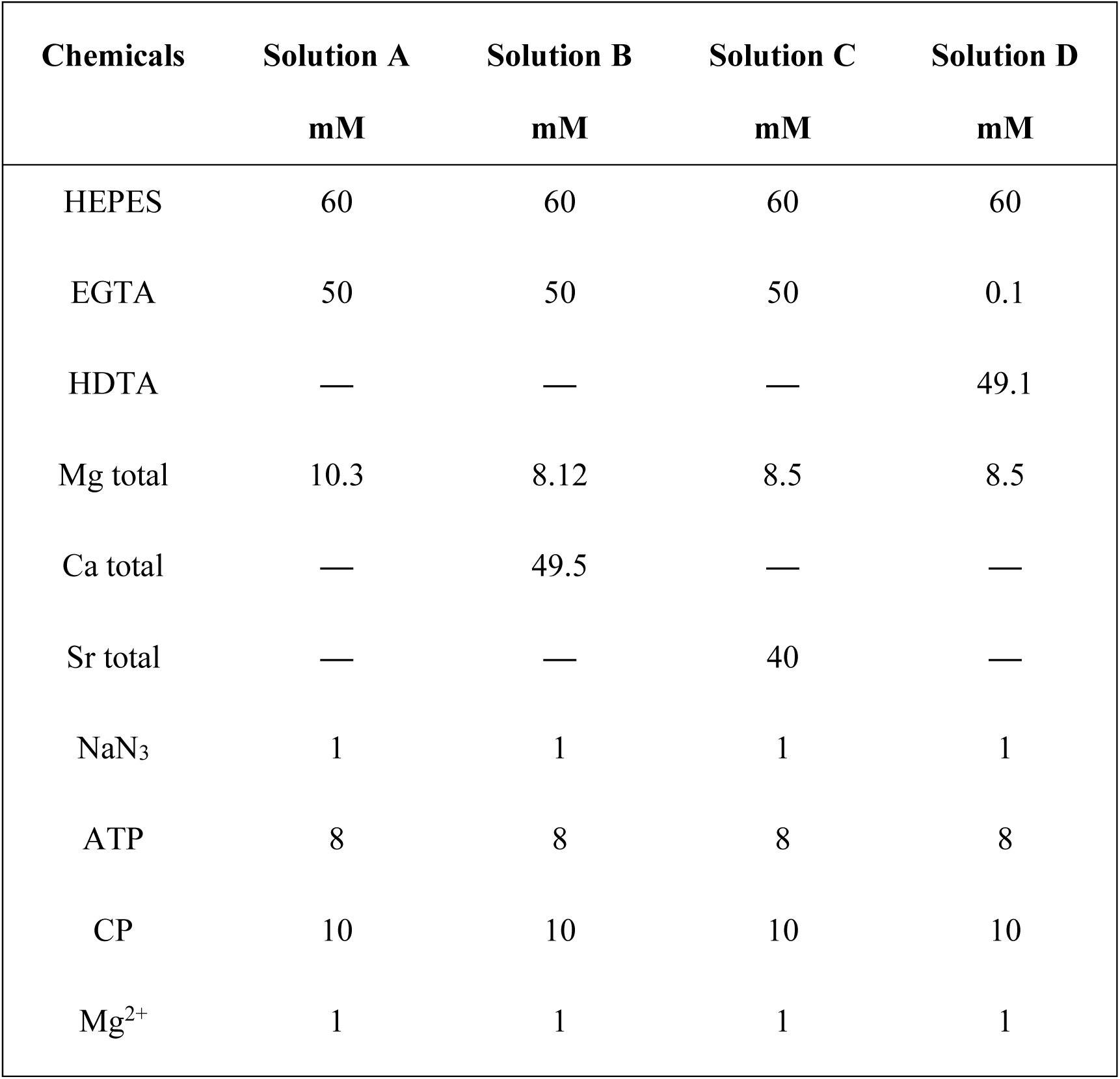
Chemical composition of stock solutions of relaxing solution (Solution A), calcium activating solution (Solution B), strontium activating solution (Solution C) and a preactivating solution (Solution D). All solutions contained (mM): K^+^ 117, Na^+^36. The pH was 7.10±0.01 at 22°C in all solutions.

| Chemicals | Solution A | Solution B | Solution C | Solution D |
| --- | --- | --- | --- | --- |
|  | mM | mM | mM | mM |
| HEPES | 60 | 60 | 60 | 60 |
| EGTA | 50 | 50 | 50 | 0.1 |
| HDTA | — | — | — | 49.1 |
| Mg total | 10.3 | 8.12 | 8.5 | 8.5 |
| Ca total | — | 49.5 | — | — |
| Sr total | — | — | 40 | — |
| NaN<sub>3</sub> | 1 | 1 | 1 | 1 |
| ATP | 8 | 8 | 8 | 8 |
| CP | 10 | 10 | 10 | 10 |
| Mg<sup>2+</sup> | 1 | 1 | 1 | 1 |

**Figure 1.**
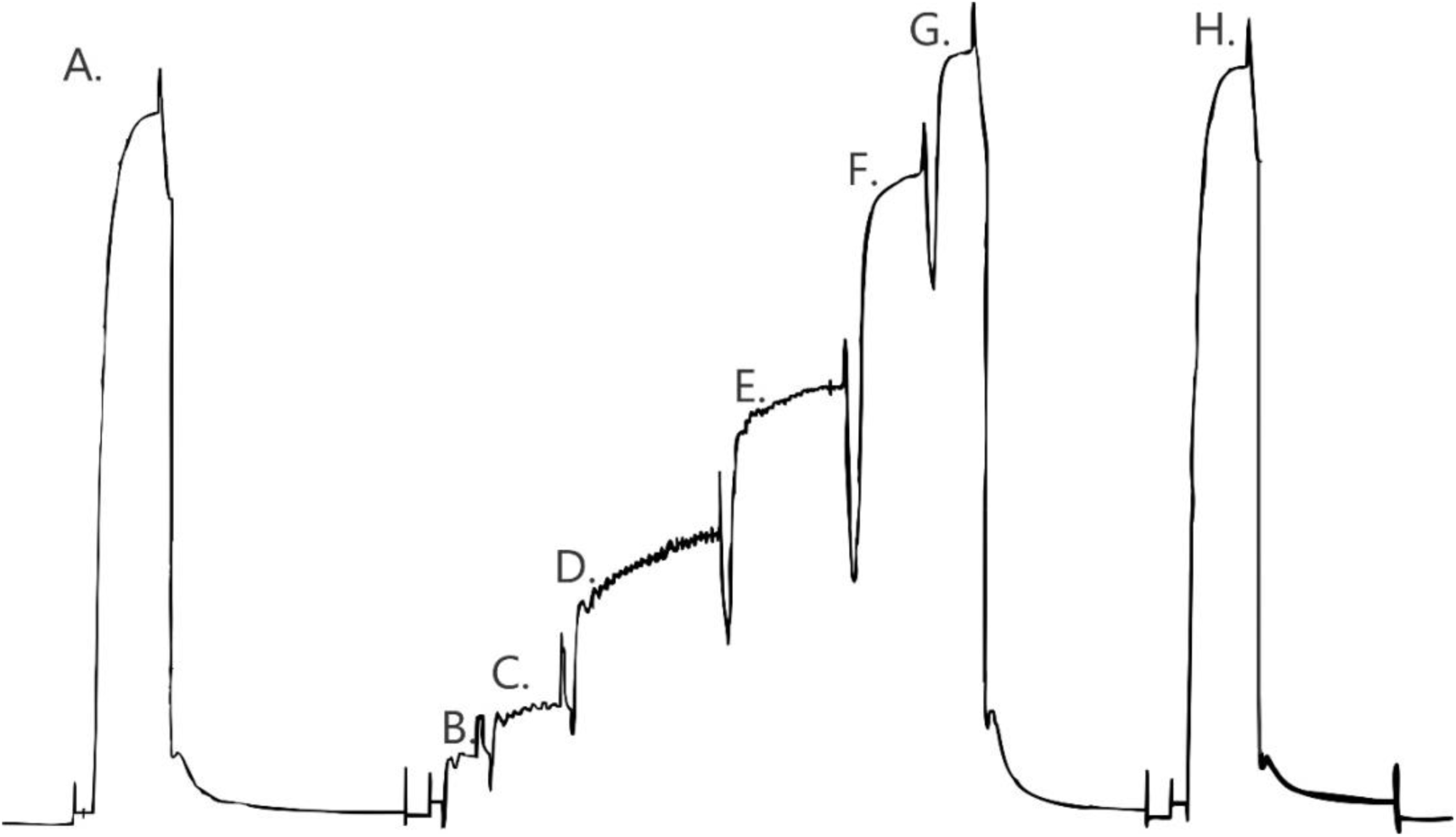
Sequence of force responses of a single skinned muscle fibre from an elderly woman, in activating solutions of increasing calcium concentrations. A, G and H: maximum force responses. B to F: force responses in solutions of increasing calcium concentrations. C,D and E: note the presence of spontaneous force oscillations of myofibrillar origin (SOMO) at these submaximal levels of force activation (frequency 0.33Hz). Duration of force activation in A is 30 seconds.

### Activation properties

Assessment of the effects of Ca^2+^ and Sr^2+^ activation was achieved by construction of force-pCa and force-pSr curves for the contractile responses of each individual fibre. The steady state tension developed in each solution was expressed as a percentage of the maximum tension developed in the sequence (Figure 1). The curves were thus generated using a modified form of the Hill equation using GraphPad Prism Software (GraphPad Software Inc., 5755 Oberlin Dr # 110 San Diego, Ca 92121). The modified Hill equation is: Relative tension (%) = 100/(1+([Ca_50_]/[Ca^2+^])^n^) where n is the Hill co-efficient for Ca^2+^ and [Ca_50_] is the calcium concentration required for half-maximal tension activation. The same equation was used for the Sr^2+^ activation curves. The following activation properties were measured from the force activation curves generated for each muscle fibre from the two age groups, when all data points fell within 5% of the fitted curves: Ca^2+^ and Sr^2+^ threshold for contraction (pCa_10_and pSr_10_, corresponding to 10% maximum force), sensitivity to Ca^2+^ and Sr^2+^ (pCa_50_ and pSr_50_, corresponding to 50% maximum force) and related differential sensitivity (pCa_50_-pSr_50_) and steepness of the activation curves (Hill co-efficient: n_Ca_and n_Sr_).

As described by Bortolotto et al. 2000, the Sr^2+^- data points for some fibres could not be well fitted by simple Hill-curves (i.e. not all data points fell within 5% of the best fitted Hill-curve). In such instances, the Sr^2+^- data points were well fitted by a biphasic curve generated by the following equation: Relative tension (%) = α/(1+([Sr_501_]/[Sr^2+^])^n1^ + β/(1+([Sr_502_]/[Sr^2+^])^n2^, where α and β represent the percentage of the two phases (α + β = 100%) and Sr_501_, Sr_502_are the strontium concentrations corresponding to the half-maximal activation of the two phases.

### Fibre classification

The Sr^2+^- dependent activation properties of individual muscle fibres permits unequivocal classification of fibres in three groups: type I (slow-twitch expressing TnC slow (cardiac) isoform and MHC I), type II (fast-twitch, expressing TnC fast isoform and MHC II isoforms) and hybrid (type I/ type II, expressing TnC fast/TnC slow and MHC I/MHC II isoforms) (O’Connell et al., 2004; Lamboley et al., 2013). This classification is based on the much higher force sensitivity to Sr^2+^ of fibres expressing the TnC slow isoform (and MHC I), than the TnC fast isoform (and MHC II isoforms) (Bortolotto et al., 2000; O’Connell et al., 2004; Lamboley et al., 2013). The hybrid fibres are characterised by biphasic force-pSr curves and express both the slow and fast TnC isoforms and a combination of MHC I and II isoforms (Bortolotto et al., 2000, O’Connell et al., 2004). Moreover, the Sr^+^-activation curves of hybrid fibres permit direct estimation of the fraction of TnC isoforms (MHC I/MHC II isoforms) present in the fibre (O’Connell et al., 2004) from the percentage ratio (α/β) of the two phases of the Sr^2+^- activation curve.

### Force oscillations of myofibrillar origin

All slow-twitch (type I) fibres and some fast-twitch (type II) and hybrid fibres displayed spontaneous force oscillations of myofibrillar origin (FOMO) at submaximal levels of force activation. The highly Ca^2+^- and Sr^2+^- buffered activation solutions used in this study (containing 50 mM EGTA) eliminates the possibility that the force oscillations were in any way caused by oscillations in Ca^2+^ or Sr^2+^ concentration. Indeed in previous studies, we have shown that these oscillations are maintained even after treatment of the fibres with detergent to disrupt and extract all membrane compartments (Stephenson & Williams, 1981). Moreover, we have shown that such force oscillations at submaximal levels of activation are caused by myosin interactions with the actin filaments (Smith & Stephenson, 2009).

### Maximum Ca^2+^ activated specific force

Skinned muscle fibres swell when exposed to relaxing solutions and the amount of swelling depends on the sarcomere length, being larger at longer sarcomeres. Therefore, when measuring the maximum Ca^2+^-activated specific force it makes sense to express the maximum Ca^2+^-activated force developed by the fibre per unit cross-sectional area before the fibre swells. The maximum Ca^2+^ activated specific force was, therefore, calculated only for fibres where the fibre diameter was measured in oil, as described by Fink et al., (1990) after mechanical skinning and before the skinned fibre was exposed to solutions. The fibre cross-sectional area was calculated assuming it to be circular.

### Statistics

The data analysed were from muscle biopsies obtained from 7 elderly women. In total, 28 muscle fibre segments were examined. Results were analysed with a one-way analysis of variance (ANOVA, Hewlett Packard statistical program) and/or Student t-test where appropriate, according to levels of significance. The Mann Whitney rank order test was applied when the sample size was small (n < 6) and the variance quite marked (SEM > 4% mean; Witte, 1993). Results were considered to be significant when P≈0.05.

## RESULTS

### Muscle fibre populations

Three populations of skeletal muscle fibres were identified from the muscle biopsies obtained from the elderly women as described in the Methods: type 1 (slow-twitch), type II (fast-twitch) and hybrid (type I/II). The slow-twitch fibres were 10 times more sensitive to Sr^2+^ than the fast-twitch fibres (pSr_50_ 5.71 vs 4.71; see Tables 2 and 3) and their identification was clear-cut. The percentages of the total muscle fibre population were: type 1 (slow-twitch) = 46%, type 2 (fast twitch) = 36% and hybrid (typeI/II) = 18%.

**Table 2.** Activation properties in slow-twitch (type I) skinned muscle fibres from elderly women circa 70 years old and older than 80 years. Mean ± standard error of the mean for (n) fibres. No statistically significant difference (P>0.05) was observed between activation properties in slow-twitch (type I) skinned muscle fibres from the two age groups of elderly women.

| **Contractile properties** | **Women circa 70 yr** | **Women > 80 yr** |
| --- | --- | --- |
| pCa<sub>10</sub> | 6.52 ± 0.05<br>(6) | 6.44 ± 0.05<br>(7) |
| pCa<sub>50</sub> | 6.12 ± 0.04<br>(6) | 6.05 ± 0.04<br>(7) |
| pSr<sub>50</sub> | 5.73 ± 0.04<br>(6) | 5.71 ± 0.06<br>(7) |
| pCa<sub>50</sub>-pSr<sub>50</sub> | 0.39 ± 0.02<br>(6) | 0.38 ± 0.04<br>(7) |
| nCa | 2.28 ± 0.13<br>(6) | 1.96 ± 0.19<br>(7) |
| nSr | 2.45 ± 0.12<br>(6) | 2.09 ± 0.26<br>(7) |

**Table 3.** Activation properties in fast-twitch (type II) skinned muscle fibres from circa 70 years old women. Mean ± standard error of the mean for (n) fibres.

| Contractile properties | Women circa 70 yr |
| --- | --- |
| pCa<sub>10</sub> | 6.29 ± 0.03<br>(10) |
| pCa<sub>50</sub> | 6.05 ± 0.02<br>(10) |
| pSr<sub>50</sub> | 4.71 ± 0.02<br>(10) |
| pCa<sub>50</sub>-pSr<sub>50</sub> | 1.35 ± 0.02<br>(10) |
| nCa | 3.38 ± 0.14<br>(10) |
| nSr | 3.33 ± 0.13<br>(10) |

When the data were divided into two age groups: fibres from elderly women circa 70 years (66 -72 yrs) and fibres from women older than 80 years, all fibres investigated from women older than 80 yrs were of type I (100%), while all three fibre types were present in women of circa 70 yrs (28.5% type I, 47.5% type II and 24% hybrid (type I/II) fibres.

### Ca^2+^- and Sr^2+^- activation characteristics

Representative force-pCa and force-pSr activation curves for slow-twitch (type I) fibres from the two age groups (circa 70 yrs and >80 yrs) of women together with the activation curves of a representative fast-twitch (type II) fibre from a 70 yrs old woman are shown in Figure 2. As described in the Methods, the following parameters were measured from the activation curves: the Ca^2+^ and Sr^2+^ thresholds for contraction (pCa_10_and pSr_10_, corresponding to 10% maximum force), sensitivity to Ca^2+^ and Sr^2+^ (pCa_50_ and pSr_50_, corresponding to 50% maximum force), related differential sensitivity (pCa_50_-pSr_50_) and steepness of the activation curves (Hill co-efficient: n_Ca_and n_Sr_). As shown in Figure 2, the Sr^2+^- activation curves lie much closer to the respective Ca^2+^-activation curves in slow-twitch fibres compared with the fast-twitch fibres.

**Figure 2.**
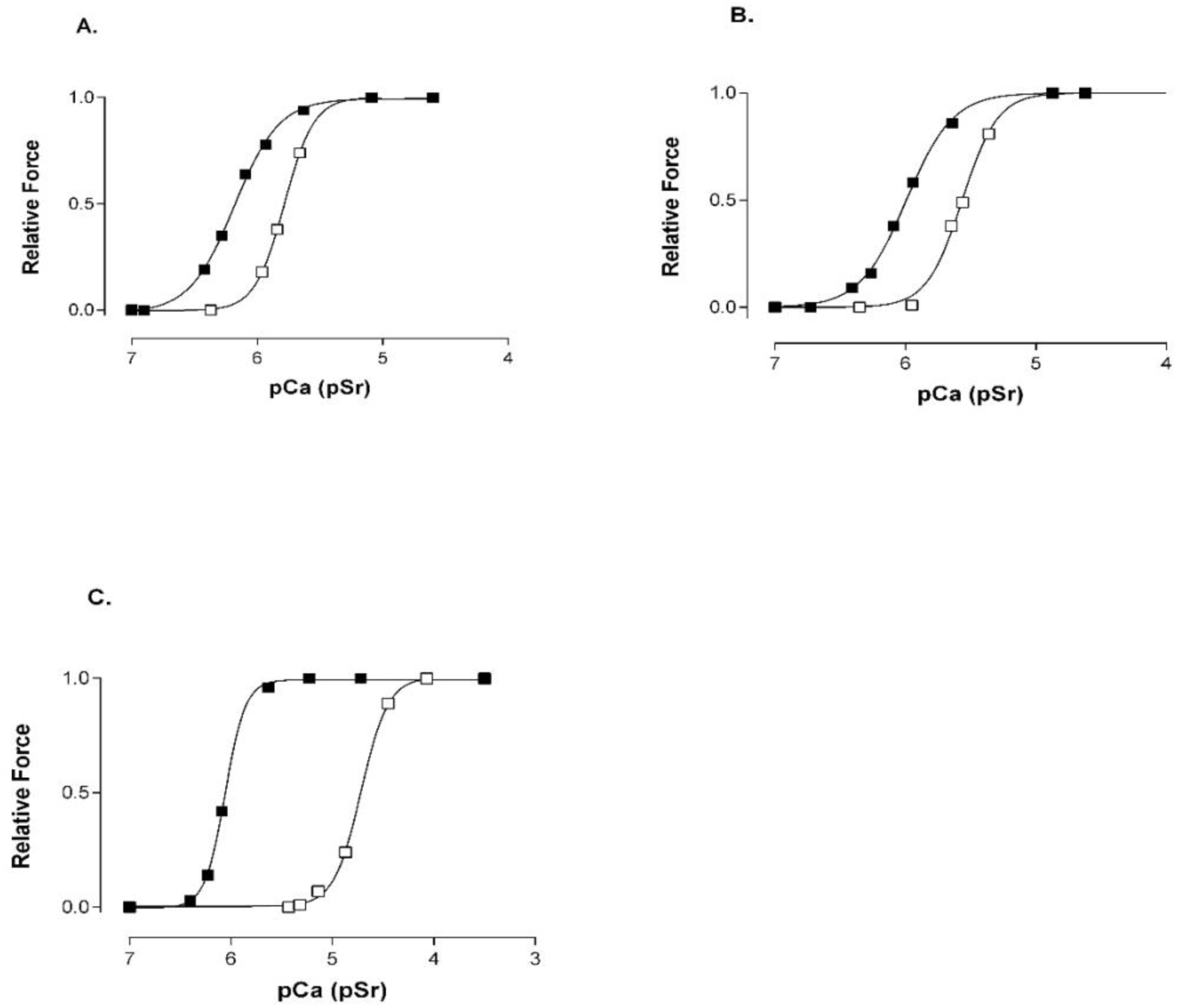
Representative force-pCa (closed symbols) and force-pSr (open symbols) activation curves in skinned muscle fibres from elderly women. A. Slow-twitch (type I) muscle fibre from a 72-year-old woman, B. Slow-twitch (type I) muscle fibre from a 90-year-old woman, C. Fast-twitch (type II) muscle fibre from a 70-year-old woman.

#### Slow-twitch (type I) fibres

The Ca^2+^- and Sr^2+^- activation characteristics of slow-twitch (type I) fibres from the two age groups (circa 70 yrs and >80 yrs) are summarized in Table 2. There were no significant differences between the measured Ca^2+^- and Sr^2+^- activation parameters of type I fibres from the two groups. However, as we point out in the Discussion, the slow-twitch (type I) fibres from older women appear to be significantly more sensitive to both Ca^2+^ and Sr^2+^, compared with the slow-twitch fibres from young adults.

#### Fast-twitch (type II) fibres

In Table 3 are shown the Ca^2+^- and Sr^2+^- activation characteristics of fast-twitch (type II) fibres from the circa 70 yrs old group. There were no type II (fast-twitch) fibres sampled from the muscle biopsies of women older than 80 yrs. The fast-twitch fibres from the circa 70 yrs old women have significantly higher Sr^2+^ and Ca^2+^ thresholds for contraction (lower pSr_10_, pCa_10_values), lower sensitivity to Sr^2+^ (lower pSr_50_ values), larger differential sensitivity (pCa_50_-pSr_50_) and higher Hill coefficients (n_Ca_, n_Sr_) than the slow-twitch fibres from the same group of women. The sensitivity to Ca^2+^ and Sr^2+^ of fast-twitch (type II) fibres sampled from the women close to 70 years of age appear to be higher than in the same type of fibres from young adults as we point out in the Discussion.

#### Hybrid fibres (type I/II)

The hybrid fibres were distinguished by their biphasic Sr^2+^- activation curve as mentioned in the Methods section. Representative Sr^2+^- and Ca^2+^-activation curves of hybrid fibres are shown in Figure 3. All fibres identified as hybrid in this study were found in the younger cohort of elderly women (circa 70 yrs). No such fibres were sampled from the biopsies of women older than 80 yrs. As shown by O’Connell et al. (2000), the ratio between the amplitudes α and β of the two phases reflects the percentage ratio between slow and fast TnC isoforms (≈ ratio of MHC I and MHC II isoforms ≈ ratio typeI/type II ≈ ratio slow-twitch/fast-twitch) expressed in the respective fibre. The percentage ratio of type I /type II myofibrillar components in the hybrid fibres identified in this study varied between 10%/90% and 50%/50%.

**Figure 3.**
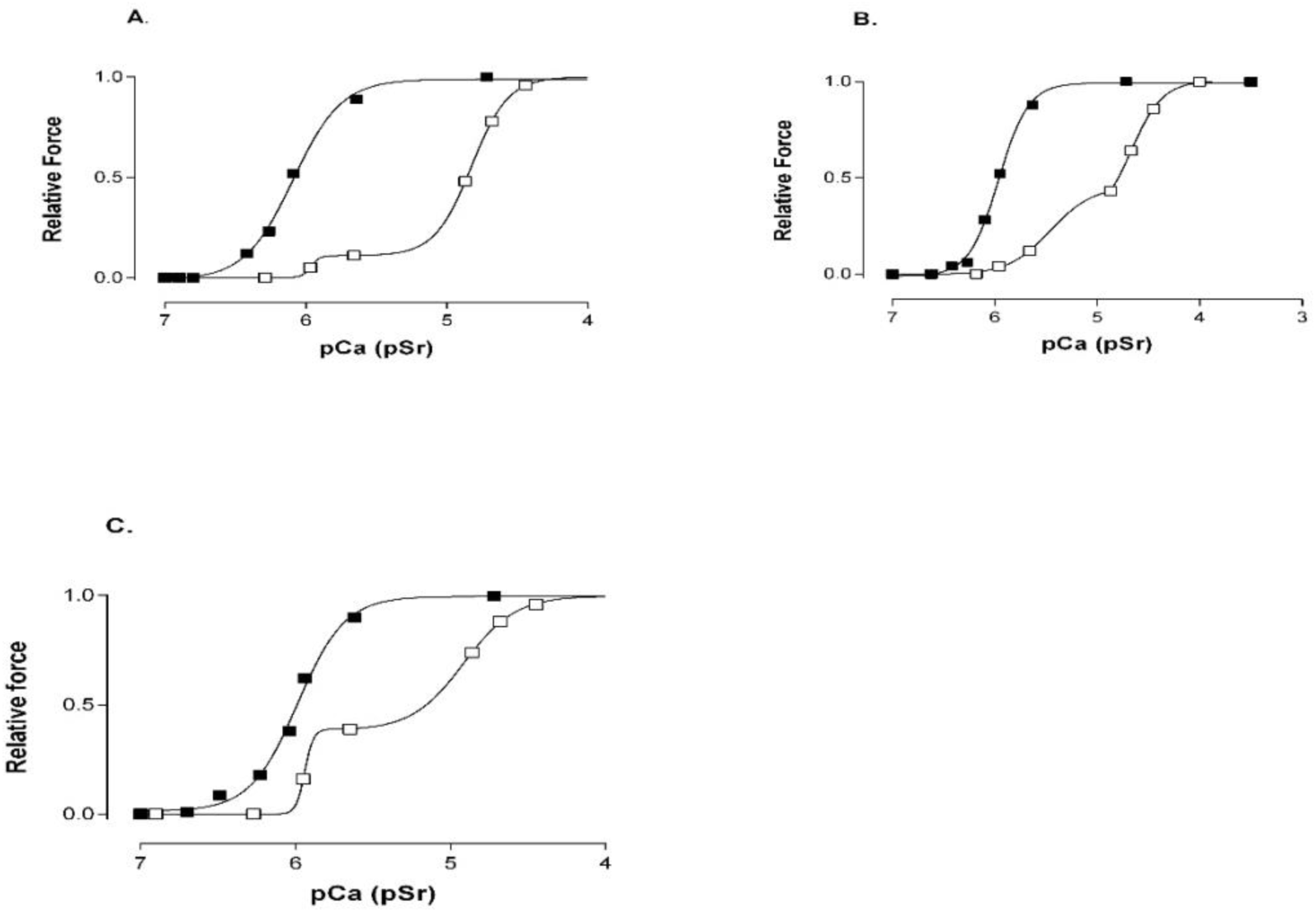
Representative force-pCa (closed symbols) and force-pSr (open symbols) curves and ratio of slow to fast twitch force activation properties in hybrid (type I/II) muscle fibres from elderly women. A. muscle fibre from a 66-year-old woman (ratio α/β = 10%/:90%). B. muscle fibre from a 66-year-old woman (ratio α/β = 50%/50%). C. muscle fibre from a 69-year-old woman (ratio α/β = 40%:/60%).

### Maximum Ca^2+^ Activated Specific Force

#### Slow-twitch (type I) fibres

The average value for the maximum Ca^2+^ activated specific force was lower in type I fibres from older women (>80 yrs) (18.3 ± 8.4 N/cm^2^, n=4) than from the women under 70 years of age (26.7 ± 8.0 N/cm^2^, n=4), but statistically, the results are not significantly different.

#### Fast-twitch (type II) fibres

There was no statistically significant difference between maximum Ca^2+^ activated specific force in fast-twitch (type 2) skinned muscle fibres from the circa 70 year old women (23.1 ± 5.7 N/cm^2^, n=4) and the slow-twitch (type I) fibres from either group of women.

#### Hybrid fibres (type I/II)

The maximum Ca^2+^ activated specific force in intermediate muscle fibres, found only in the circa 70 yrs old group of women, varied between 16.3 to 40.0 N/cm^2^ and was not statistically different from the values of either the slow- or the fast-twitch fibres in the two groups of women.

#### Pooled force data from all fibres

As detailed in the Discussion, the maximum Ca^2+^ activated specific force in the pooled data from this study was significantly lower by about 20% than the pooled data for young adults from a study from the same laboratory using similar solutions.

### Frequency of Force Oscillations of Myofibrillar Origin (FOMO)

#### Slow-twitch (type I) fibres

Figure 4 shows representative force responses in submaximally activated fibres where the FOMO phenomenon was observed and the results are summarized in Table 4. In slow-twitch (type I) skinned muscle fibres the frequency of FOMO was not significantly different between the two age groups of elderly women. The FOMO phenomenon was very regular in the slow-twitch (type I) fibres from the circa 70 year old women, but showed irregularities in some slow-twitch fibres from women > 80 years (Figure 4 B, C).

**Table 4.** Frequency (Hz) of force oscillations of myofibrillar origin (SOMO) in slow-twitch (type I), fast-twitch (type II) and hybrid (type I/II) skinned muscle fibres. Mean ± standard error of the mean (n = force traces showing SOMO). **\*** Significant difference (P≤0.02) between frequency of SOMO in fast-twitch (type II) skinned muscle fibres from circa 70 years old women and those from adults (Fink et al., 1990).

| Age | Slow-twitch | Fast-twitch | Hybrid |
| --- | --- | --- | --- |
| c. 70 yr old<br>women | 0.15 ± 0.01<br>(11)<br>regular | 0.21 ± 0.07 *<br>(3)<br>irregular | 0.17 ± 0.03<br>(3)<br>irregular |
| > 80 yr old<br>women | 0.19 ± 0.06<br>(4)<br>slightly irregular | No fast-twitch<br>fibres | No hybrid<br>fibres |
| Adults | 0.15 ± 0.01<br>(7)<br>regular | 0.5 ± 0.07 *<br>(7)<br>irregular | 0.18 and 0.23<br>(2) |

**Figure 4.**
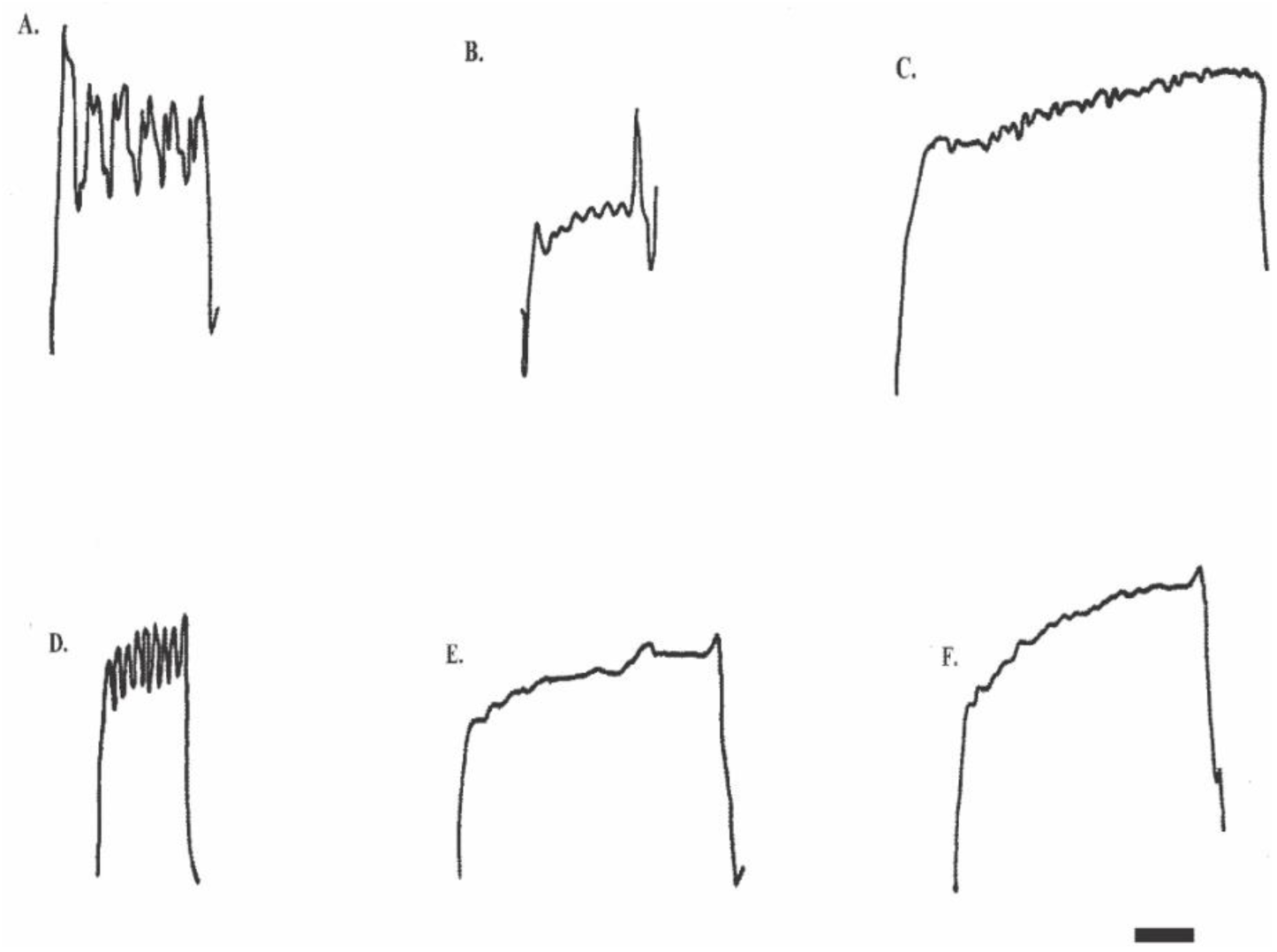
Representative tracings of force oscillations of myofibrillar origin in: A. Slow-twitch skinned muscle fibre from a 72-year-old woman (regular force oscillations 0.11 Hz). B. Slow-twitch skinned muscle fibre from a 90-year-old woman (regular force oscillations 0.22 Hz). C. Slow-twitch skinned muscle fibre from a 90 a woman (irregular force oscillation 0.11 Hz). D. Fast-twitch skinned muscle fibre from a 72-year-old woman (regular force oscillations 0.33 Hz). E. Fast-twitch skinned muscle fibre from a 72-year-old woman (irregular force oscillations 0.10 Hz). F. Hybrid (type I/II) skinned muscle fibre from a 66-year-old woman (irregular force oscillations, 0.22 Hz). Horizontal scale bar = 20 seconds in all panels.

#### Fast-twitch (type II) fibres

The FOMO phenomenon was observed in a small number of fast-twitch (type II) skinned muscle fibres from elderly women, circa 70 years old. As shown in Table 4 and Figure 4 (D, E) the force oscillations in these fibres were not always regular and had frequencies that were not significantly different from those in slow-twitch (type I fibres).

## DISCUSSION

The results obtained in this study show that with advancing age, in women, there are notable changes in the vastus lateralis muscle viewed from the perspective of fibre type ratio and single fibre contractile properties. Based on their functional properties with respect to activation by Sr^2+^ the fibres could be grouped into three distinct categories: slow-twitch (type I), fast-twitch (type II) and hybrid (type I/II) fibres. All three fibre types were present in the biopsies of women of circa 70 yrs of age, but no fast-twitch or hybrid fibre was sampled from biopsies of women older than 80 yrs. This may be a consequence of taking one small sample from a specific region in the women of advanced age because slow motor units reinnervate the adjacent denervated fast-twitch muscle fibres transform them into slow motor units containing large numbers of slow muscle fibres all present in one area rather than being distributed throughout the muscle is in case younger subjects Piasecki et.al. 2016. Hybrid fibres may represent fibres in different states of transformation/degeneration/regeneration (Begam & Roche 2018, Matsuura et.al. 2007). Their relatively large proportion (24%) in the vastus lateralis muscle of women of circa 70 yrs of age and absence of such fibres in the same muscle of women older than 80 yrs may suggest that there are intense processes of muscle remodelling occurring around the seventh decade of life in women. The decline in the population of fast-twitch fibres in older animals and humans is possibly due to their increased susceptibility to damage (Aniansson et al., 1980; Larsson et al., 1978; Lexell et al., 1988; Lexell, 1995, Chan & Head 2010) and also as mentioned above due to preferential loss of fast motor units as a consequence of the loss of fast motor neurons during advanced ageing (Piasecki et.al. 2016).

Moreover, it appears that there are distinct functional differences between fibres of the same type in women of advancing age and young adults. For example, a recent study on the vastus lateralis muscle from young adults (16 males, seven females, age 24 +/-5 yrs) (Lamboley et al., 2013) using the same activating solutions and equipment as in the present study, and a similar system for fibre type classification, showed that the pCa_50_ values for slow-twitch (type I) and fast-twitch (type II) fibres from young adults were 5.93 +/-0.01 (n=40) and 5.83 +/-0.01 (n=30) respectively. In comparison, the slow- and fast-twitch fibres from the vastus lateralis muscle of elderly women (66-90 yrs) had significantly greater pCa_50_ values (6.09 +/-0.03, n=13 for slow-twitch fibres and 6.05 +/-0.02, n =10 for fast-twitch fibres). On average, the results show that muscle fibres from elderly women produce 50 % of their maximum Ca^2+^-activated force at Ca^2+^ concentrations that are lower by a factor of 1.4 for slow-twitch fibres and 1.7 for fast-twitch fibres compared with fibres of the same type from young adults. The Ca^2+^-activation curves of both slow- and fast-twitch fibres also appear to be significantly less steep in fibres from elderly women than in fibres from young adults as indicated by the smaller Hill-coefficients in the elderly women (n_Ca_= 2.11 +/-0.15 vs 5.2 +/-0.2 for slow-twitch and 3.38 +/-0.14 vs 5.1+/-0.1 for fast-twitch fibres).

The percentage of hybrid fibres (type I/II) appears to be several-fold higher in the 66-72 yrs group of elderly women (24%) than in adults (< 5%, Lamboley et al., 2013; <10% Fink et al., 1990), further supporting the suggestion that muscle remodelling may occur around the seventh decade of life in women. A proportion of this remodeling is likely a consequence of the preferential loss of fast twitch-motor neurons with advancing age Lexell 1995, Lexell & Taylor 1991.

The average maximum Ca^2+^-activated specific force expressed per cross sectional area of the skinned fibre before swelling, measured in this study, was about 20% lower in the pooled fibres from all elderly women (24.3 N/cm^2^) compared with the average value for adult fibres from the vastus lateralis (30.0 N/cm^2^) obtained in a previous study from the same laboratory using similar procedures and solutions, but where the solutions contained, in addition, 10 mM caffeine (Fink et al., 1990). In a recent review on measuring specific force in human single chemically skinned muscle fibres Kalakoutis and colleagues (Kalakoutis et al. 2021) reported a five-fold differences in mean specific force data reported from different laboratories, however, importantly for the comparison we are making here with our earlier study Fink et al. 1990, they observed a consisted specific force reported for research groups from the same laboratory using similar technique and solutions. Note, however, that there was no difference between the maximum Ca^2+^ activated specific force in chemically skinned fibres of the same type from young and elderly women, when the force measurements were made at 15°C (Trappe et al., 2003). Nevertheless, in their study, Trappe et al., (2003) found that under their conditions, the maximum specific force was about 40% greater in fast-twitch than in slow-twitch fibres. Considering the decline in the number of fast-twitch fibres in the vastus lateralis muscle of elderly women, this would translate to lower overall forces per cross-sectional area in elderly women. Miller et al. 2013 showed that aging slows myosin actin cross-bridge kinetics in women, leading to decrements in whole body dynamic contractile performance.

The spontaneous force oscillations that originate from the interaction of myofibrillar proteins (SOMO, see Figure 4 and Table 4) when skinned fibre are submaximally activated is characterised by the frequency of the oscillations, which is fibre type specific, and their regularity (Smith & Stephenson 1994) furthermore, the presence of these oscillations is an indication of the health of the single fibres obtained from the human biopsy demonstrating the contractile proteins have not deteriorated either in the sampling process or for the five hours which they were used in the laboratory. The frequency of SOMO in fast-twitch (type II) skinned muscle fibres from circa 70 year old women was significantly different, approximately half of that in the equivalent fibre type from adults (Fink et al., 1990). In general, SOMO in the fast-twitch (type II) fibres of the elderly women was more similar to SOMO observed in the hybrid (type I/II) fibres from adults than in the fast-twitch (type II) fibres of adults (Fink et al., 1990), further suggesting that the contractile proteins of the fast-twitch (type II) muscle fibres may have been altered in some way by the aging process.

Taken together these results show that with advancing age, in women, there are changes in the activation properties of the contractile apparatus of muscle fibres from the vastus lateralis muscle. There is also an increase in the number of slow fibres and hybrid fibres and a decrease in fast-twitch fibres. These observed changes occur on the background of a robust capacity of old and senescent muscle to regenerate functional architecture (Lee et al. 2013) and need to be taken into consideration when explaining observed changes in women’s mobility with aging.

